# Optimizing ex vivo CAR-T cell-mediated cytotoxicity assay through multimodality imaging

**DOI:** 10.1101/2024.05.03.592348

**Authors:** John G. Foulke, Luping Chen, Hyeyoun Chang, Catherine E. McManus, Fang Tian, Zhizhan Gu

## Abstract

CAR-T cell-based therapies have demonstrated remarkable efficacy in treating malignant cancers, especially liquid tumors, and are increasingly being evaluated in clinical trials for solid tumors. With the FDA’s initiative for advancing alternative methods for drug discovery and development, full human ex vivo assays are increasingly essential for precision CAR-T development. However, prevailing ex vivo CAR-T cell-mediated cytotoxicity assays are limited by their use of radioactive materials, lack of real-time measurement, low throughput, and inability to automate, among others. To address these limitations, we optimized the assay using multimodality imaging methods, including bioluminescence, impedance tracking, phase contrast, and fluorescence, to track CAR-T cells co-cultured with CD19, CD20, and HER2 luciferase reporter cancer cells in real-time. Additionally, we varied the ratio of CAR-T cells to cancer cells to determine optimal cytotoxicity readouts. Our findings demonstrated that the CAR-T cell group effectively attacked cancer cells, and the optimized assay provided superior temporal and spatial precision measurements of ex vivo CAR-T killing of cancer cells, confirming the reliability, consistency, and high throughput of the optimized assay.

## Introduction

CAR-T therapy, also known as chimeric antigen receptor T-cell therapy, is an innovative and promising form of immunotherapy that has revolutionized the field of cancer treatment (June et al., 2018). This cutting-edge therapy involves reprogramming a patient’s own T cells to express a chimeric antigen receptor (CAR) specific to tumor-associated antigens (TAAs) (Sadelain et al., 2013). By genetically modifying T cells to recognize and target cancer cells with precision, CAR-T therapy offers a personalized and targeted approach to treat various types of hematological malignancies and solid tumors. The remarkable success of CAR-T therapy in clinical trials and its approval for certain indications such as leukemias, lymphomas, and multiple myelomas, have marked a significant breakthrough in cancer treatment (Garfall et al., 2015; Kochenderfer et al., 2010; Porter et al., 2011; Sengsayadeth et al., 2022). As ongoing research continues to refine CAR-T cell design, improve safety profiles, and expand the range of treatable cancers, this transformative therapy holds the potential to reshape the landscape of oncology and pave the way for more effective and personalized cancer therapies.

Considerable research efforts have been invested into developing new CAR structures to increase the scope of targeted cancer types and raise their anti-tumor efficacy (Jackson et al., 2016). One of the bottlenecks in the process of CAR-T development is evaluating the biofunction of CAR-T cells ex vivo. Despite the growing importance of ex vivo CAR-T cell cytotoxicity assays as a valuable means of assessing CAR-T cell functionality and effectiveness, there are still various challenges and limitations that need to be addressed. These include simplified tumor microenvironment, limited immune system interactions, artificial activation and expansion, lack of dynamic monitoring, inadequate assessment of CAR-T cell exhaustion and variability in assay techniques (Kiesgen et al., 2021).

As recently reviewed on Nature Protocols, the prevailing ex vivo CAR-T cytotoxicity assays are the chromium (^51^Cr)–release assay (^51^Cr assay), the luciferase-mediated bioluminescence imaging (BLI) assay, the impedance-based assay, and the flow cytometry assay (flow assay) (Kiesgen et al., 2021). However, each of these assays has its own limitations. For example, the chromium release assay is a commonly used method to measure target cell lysis by CAR-T cells. It involves labeling target cells with a radioactive chromium isotope, which is released into the supernatant upon target cell lysis (Brunner et al., 1968). While widely used, the limitations of this assay include radioactivity, lack of dynamic monitoring, non-specific release, insensitivity to low levels of killing and delayed measurements (Erskine et al., 2012; Peper et al., 2014; Riss et al., 2019). The flow cytometry assay is a widely used technique for evaluating CAR-T cell cytotoxicity by measuring target cell death or marker expression. However, it also has some limitations such as limited dynamic range, lack of spatial information, influence of non-specific binding and limitations of fluorophores. The impedance-based assay is limited by its lack of specificity, insufficient sensitivity, difficulty in multiplexing, indirect measurement, and insensitivity to low-level cytotoxicity, etc. The limitations of the bioluminescence imaging assay commonly include signal attenuation, lack of spatial information, signal persistence, background noise, and limited resolution (Karimi et al., 2014; Kiesgen et al., 2021; Riss et al., 2019).

Overall, every single ex vivo CAR-T cell-mediated cytotoxicity assay has various limitations due to the assay technology itself. We generated a panel of luciferase reporter tumor cell lines that can be used to examine the function of CAR-T cells. The reporter cells in this panel naturally express high levels of clinically relevant CAR-T target antigens on cell surface, such as CD19, CD20 and HER2. By utilizing stable luciferase expression in these CAR-T target tumor cell lines, we can enhance the ex vivo CAR-T cell-mediated cytotoxicity assay by employing multimodality imaging techniques. This approach allows us to overcome certain limitations associated with individual single assay methods, such as radioactivity and the absence of spatial information, while optimizing overall assay performance.

## Results

### mRNA profiling of cancer cell Lines for common CAR-T targets

Over 30 luciferase expressing cell lines have been generated at ATCC and are maintained in the extensive cancer cell line collection. To identify the best candidate lines for CAR-T cytotoxicity assay development, we examined expression data from the CCLE database for transcript levels of 12 common hematological tumor CAR-T targets and 26 common solid tumor CAR-T targets (Chu et al., 2018; Martinez and Moon, 2019). Data was available for the parental cell lines of 16 existing luciferase-expressing lines at ATCC representing 16 different tissue and cancer types. We created a heatmap in which genes were colored red to indicate high expression and blue to indicate low expression (Fig 1). Few cell lines with existing ATCC luciferase reporters have high levels of CD19 or BCMA, the only two FDA-approved CAR-T targets as of now (Cappell and Kochenderfer, 2023). When we examined expression data for additional ATCC parental cancer cell lines without already existing luciferase reporters, we found that the lymphoma cell lines Raji, Farage, and Daudi express high levels of CD19, as well as high levels of CD20, CD22 and CD38. In addition, BT-474, a breast ductal carcinoma cell line, expresses high levels of HER2, another very popular CAR-T target in solid tumors currently under investigation (Budi et al., 2022). WIL2-S, a B lymphoblast cell line, did not have available expression data in the CCLE database but was included due to known high levels of CD20 (Gazzano-Santoro et al., 1997). In-house RNAseq experiments confirmed high expression of CD19, CD20, CD22, and CD70, etc. in both WIL2-S and RAJI cell lines (Fig 2A). Overall, these results show that key CAR-T targets are endogenously expressed at high levels in these cell lines, making them good candidates for generating CAR-T target luciferase reporter lines. In addition, this analysis can be used to determine which luciferase-expressing cell lines available from ATCC could be good candidates for use in studies of other CAR-T targets such as LeY, ROR1, or WT1 in hematological tumors and MET, CD70, or EPCAM in solid tumors (Chu et al., 2018; Martinez and Moon, 2019).

**Figure 1:**
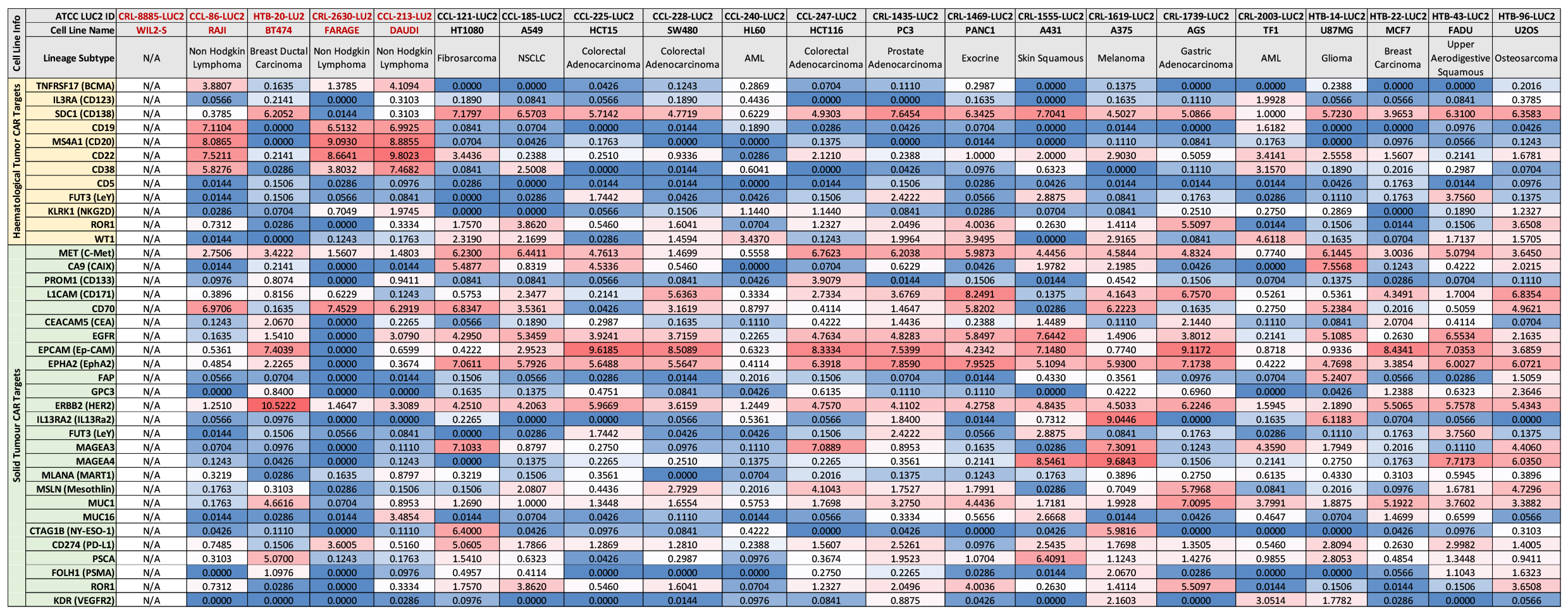
Analysis of common hematologic and solid tumor CAR-T targets in selected cancer cell lines. Heat map of expression levels from the CCLE database for 12 hematological and 26 solid tumor CAR-T targets in 21 cancer lines with luciferase-expressing daughter lines available from ATCC. Values are log2(TPM+1). Note: WIL2-S did not have RNA-seq data within the CCLE database.

**Figure 2:**
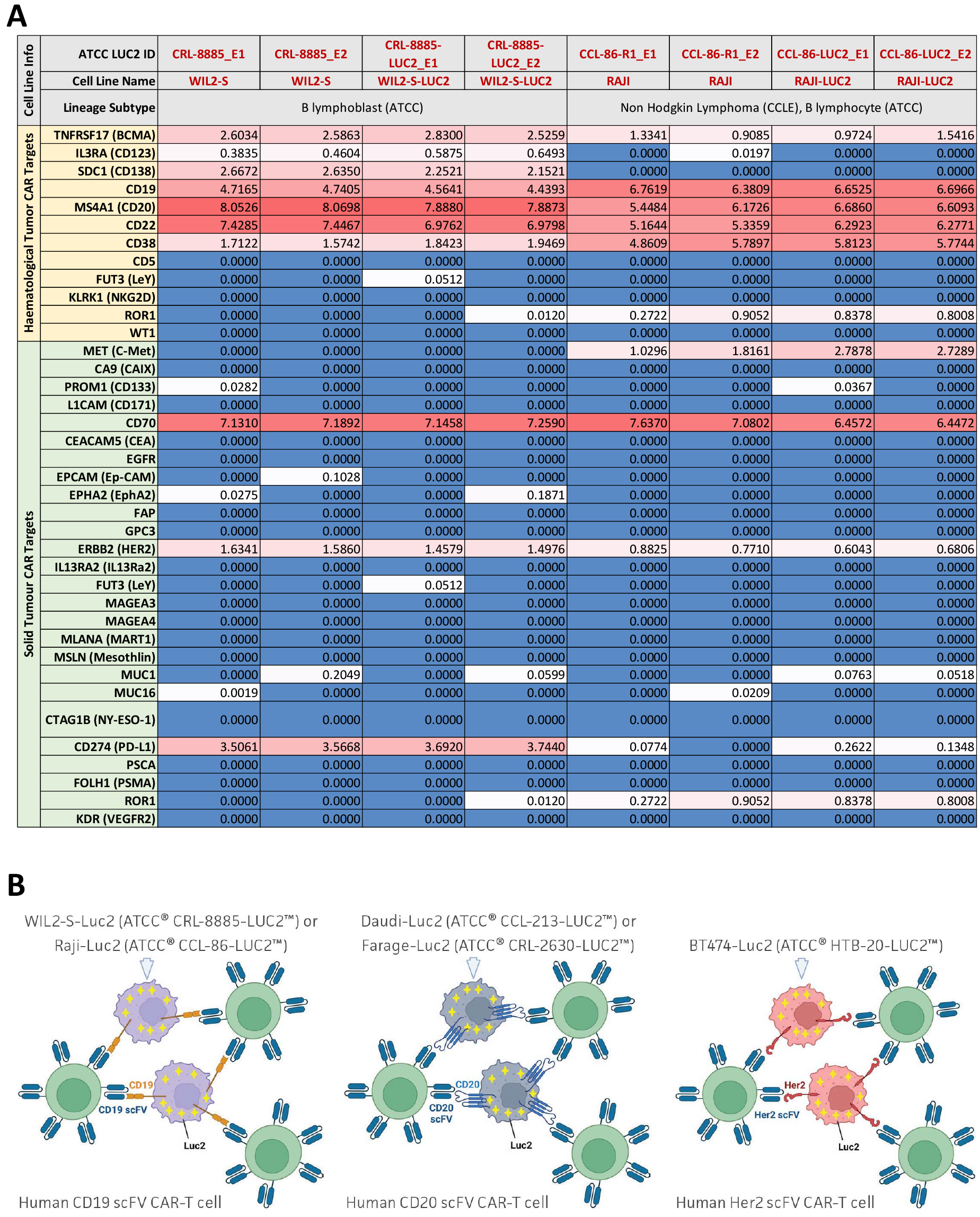
mRNA Seq analysis of common hematologic and solid tumor CAR-T targets in B lymphoblast and B lymphocyte cancer cell lines. A) Heat map of RNA-seq data for 2 cancer cell lines and 2 luciferase-expressing daughter cell lines showing expression levels of 12 hematologic tumor CAR-T targets and 26 solid tumor CAR-T targets. RNA sequencing was performed in duplicate by Psomogen. CD19 expression remains consistent pre- and post-luciferase transduction. Values are log2(FPKM+1). B) Schematic showing CD19-positive WIL2-S-Luc2 and Raji-Luc2, CD20-positive Daudi-Luc2 and Farage-Luc2, and HER2-positive BT-474-Luc2 cells being surrounded and attacked by CD19-, CD20-, and HER2-targeting CAR-T cells, respectively. Created with BioRender.com.

We therefore generated 5 new luciferase reporter cell lines for potential use in CAR-T cell therapy research: Raji-Luc2, WIL2-S-Luc2, BT-474-Luc2, Daudi-Luc2, and Farage-Luc2 (Fig 2B). A lentiviral plasmid was transduced into these ATCC authenticated tumor cell lines, and single cell cloning was performed to generate a homogenous culture which was expanded and validated for luciferase expression. Single clones with high luciferase activity were further characterized by comparing their morphology and growth kinetics to their parental cell lines, which revealed high similarity between parental and daughter Luc2 lines (Figs S1 and S2). High expression levels of endogenous CAR-T target antigens in the reporter lines were also verified by flow cytometry (Fig S3). Finally, the reporter lines showed luciferase expression that increased linearly with cell number and was stable over at least 30 generations (Figs S4 and S5).

To compare the expression levels of CAR-T target antigens in engineered versus parental lines, mRNA sequencing was performed on cell pellets isolated from WIL2-S, WIL2-S-Luc2, Raji, and Raji-Luc2, in duplicate. Results confirmed that common CAR-T target expression in the Raji and WIL2-S parental and daughter lines remains unaffected by the introduction of luciferase (Fig 2A).

### Optimizing ex vivo CD19 CAR-T cell-mediated cytotoxicity assay using luminescence and live cell imaging

To investigate whether the newly generated CAR-T luciferase reporter cell lines could be used to study the cytotoxic effects of antigen-specific CAR-T cells, we performed co-culture experiments using commercially available CAR-T cells specifically targeting CD19. CD19 CAR-T cells were paired with mock CAR-T control cells from the same donor. This allowed us to compare mock CAR-T cells and target-specific CAR-T cells in the same genetic background and with similar levels of non-specific killing activity. CD19-positive Raji-Luc2 cells or WIL2-S-Luc2 cells were co-cultured with either CD19 CAR-T or mock CAR-T cells at various ratios of CAR-T cells to target cells (1:1, 2:1, 5:1, and 10:1). After 24 hours of co-culture, cell killing was measured via luminescence of the reporter cells. When the luminescence signal was normalized to wells containing only reporter cells, luminescence decreased with increasing amounts of CAR-T cells (Fig 3A and B). Importantly, luminescence decreased significantly more when reporter cells were co-cultured with CD19 CAR-T cells compared to nonspecific levels of killing after co-culture with mock CAR-T cells. Overall, these results demonstrate how reduction in luminescence in these reporter lines can be used as a proxy for CAR-T targeting efficacy.

**Figure 3:**
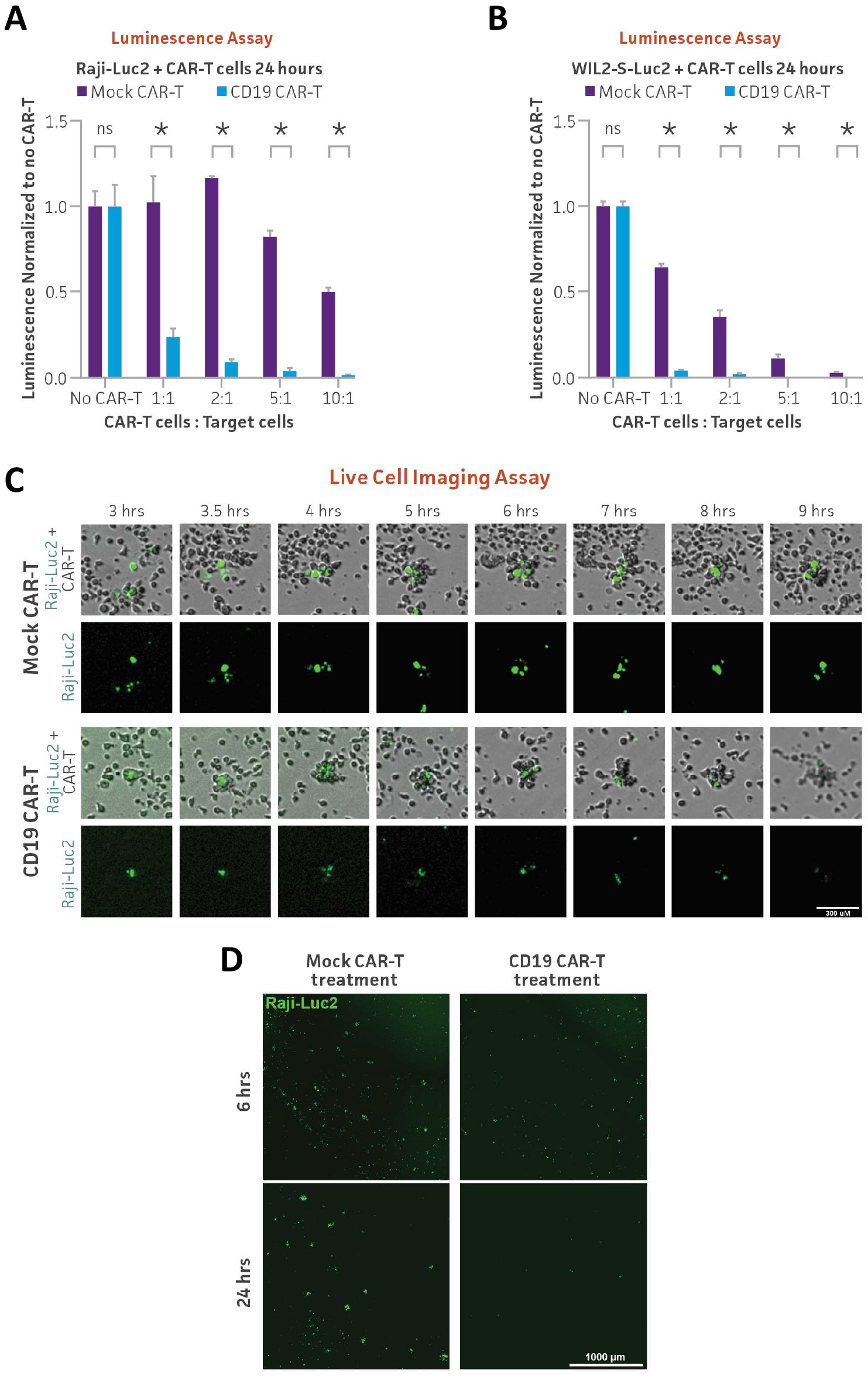
CD19 CAR-T in vitro killing assay of Raji-Luc2 and WIL2-S-Luc2 measured using luminescence and live cell imaging. (A) CD19-positive Raji-Luc2 cells (5 x 10^3^) or (B) WIL2-S-Luc2 cells (5 x 10^3^) were seeded into a 96-well plate and were used as target cells for either CD19 CAR-T or mock CAR-T cells (control) from the same donor, which were seeded at various ratios of CAR-T cells to target cells (1:1, 2:1, 5:1, and 10:1). After 24 hours of co-culture, Bright-Glo reagent was added to the wells. Luminescence was read within 10 minutes using a plate reader and was normalized to wells with no CAR-T cells. * = significant difference and ns = not significant using an unpaired t test, with a single pooled variance. N=3 in all experiments. *P < 0.05. (C) Raji-Luc2 cells were stained with Vybrant DiO dye and real-time fluorescent images were captured every 30 minutes for 24 hours during co-culture with either CD19 or mock CAR-T cells. Stained Raji-Luc2 cells (green) from the co-culture experiment were tracked for 6 hours and became surrounded by CAR-T cells, resulting in a decrease in fluorescence when treated with CD19 CAR-T cells as compared to co-cultures with mock-CAR-T cells. Scale bar, 300 μm. (D) The larger fields of view of Vybrant DiO stained Raji-Luc2 cells (green) after 6 or 24 hours of co-culture with either CD19 or mock CAR-T cells. Scale bar, 1000 μm.

While bioluminescence is a powerful and simple read-out for cytotoxicity, it is not as convenient to monitor cell killing in real time as fluorescence-based live cell imaging (Tung et al., 2016). To evaluate CAR-T cytotoxicity using a real-time assay, Raji-Luc2 cells were stained with lipophilic fluorescent DiO dye and fluorescent images were captured every 30 minutes for 24 hours during co-culture with either CD19 CAR-T cells or mock CAR-T cells. When Raji-Luc2 cells were co-cultured with mock CAR-T cells, fluorescence levels decreased slowly and remained relatively constant, even as the reporter cells became surrounded by CAR-T cells. However, a quick decrease in fluorescence, indicating the massive killing of target cells, was observed when stained Raji-Luc2 cells were co-cultured with CD19 CAR-T cells (Fig 3C; Video 1 and Video 2). Visualizing a large field of cells displayed a significant fluorescence decrease observed at 24 hours after co-culture with CD19 CAR-T cells while no significant change in fluorescence was observed after co-culture with mock CAR-T cells (Fig 3D). These results indicate that the CAR-T target reporter lines can be used to monitor CAR-T cell targeting using fluorescence-based live imaging techniques in addition to bioluminescence.

### Optimizing ex vivo HER2 CAR-T cell-mediated cytotoxicity assay using luminescence and xCELLigence live cell assay in adherent cells

Improving the efficacy of CAR-T cells on solid tumors remains an active area of research (Dagar et al., 2023). To investigate whether the BT-474-Luc2 line could be used as a reporter of HER2 CAR-T cell targeting, cytotoxicity of HER2 CAR-T cells on BT-474-Luc2 cells was measured by luminescence. HER2-positive BT-474-Luc2 cells were seeded at the same CAR-T to target cell ratios as above using either HER2 CAR-T or mock CAR-T cells. After 24 hours of co-culture, luminescence dramatically decreased with increasing doses of HER2 CAR-T cells (Fig 4A). This result demonstrates that the BT-474-Luc2 reporter line can be used to monitor HER2 CAR-T cell targeting via luminescence.

**Figure 4:**
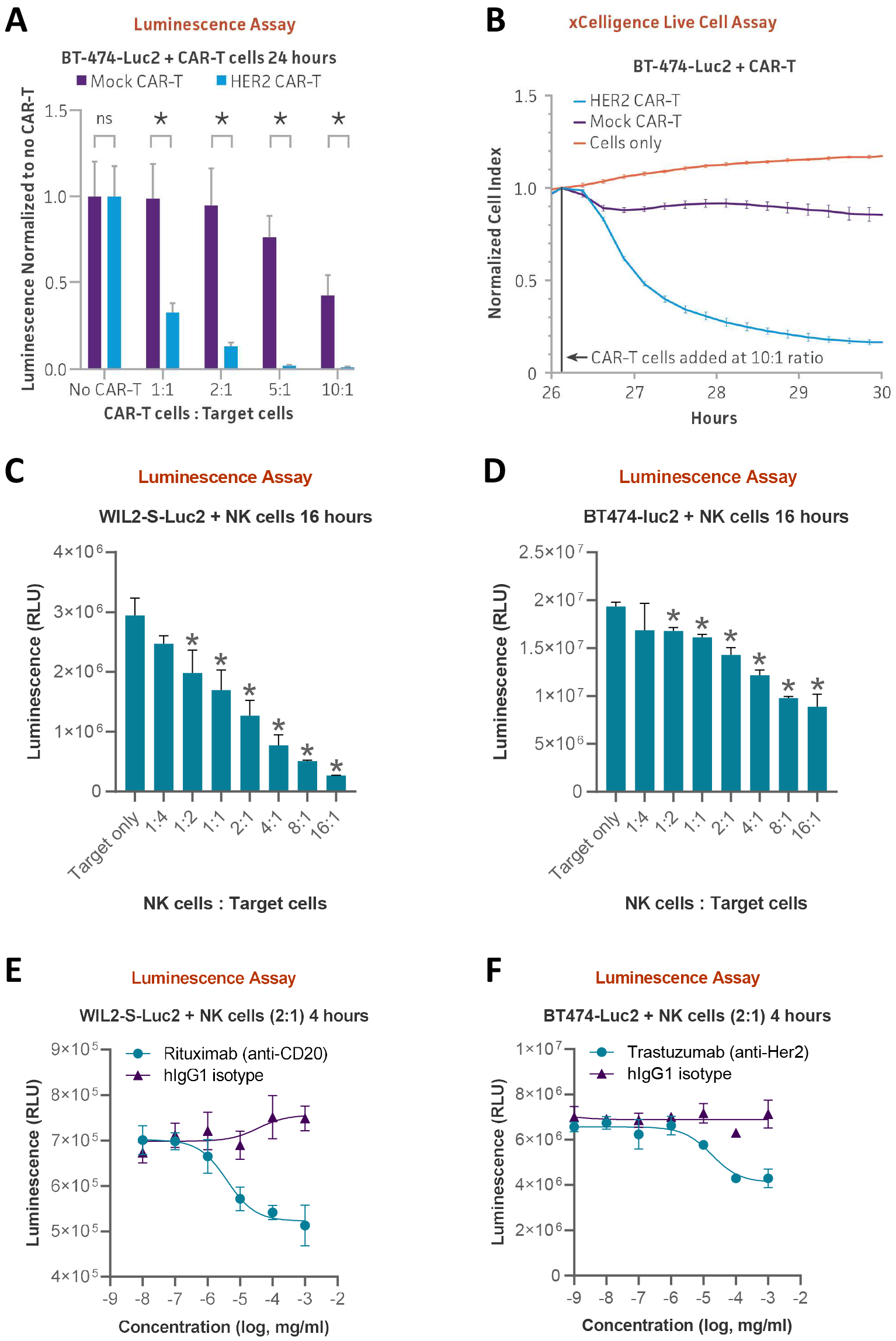
BT-474-Luc2 and WIL2-S-Luc2 used in CAR-T and NK cytotoxicity assays. (A) HER2-positive BT-474-Luc2 cells (5 x 10^3^) were seeded into a 96-well plate and either HER2 CAR-T or mock CAR-T cells were added at various ratios of CAR-T cells to target cells (1:1, 2:1, 5:1, and 10:1). (B) HER2 CAR-T cells were used to target 2 x 104 HER2-positive BT-474-Luc2 cells at a 10:1 ratio and cell killing was measured using the xCELLigence system. Mock CAR-T cells from the same donor were used as a control. (C) WIL2-S-Luc2 cells or (D) BT-474-Luc2 cells were co-cultured with NK cells for 16 hours at various NK to target cell ratios (1:4, 1:2, 1:1, 2:1, 4:1, 8:1, and 16:1) after which luciferase activity was measured. E) Antibody-dependent cell-mediated cytotoxicity (ADCC) assay using Rituximab (anti-CD20) with WILS-2-Luc2 cells or F) Trastuzumab (anti-HER2) monoclonal antibodies with BT474-Luc2 cells. Concentrations of monoclonal antibody or IgG1 control varying from 10 pg to 1 μg/mL were added to co-cultures of primary NK cells and reporter cells at a 2:1 NK:target cell ratio. Luminescence was measured after 4 hours of co-culture. * = significant difference and ns = not significant using an unpaired t test, with a single pooled variance. N=3 in all experiments. *P < 0.05.

As the only adherent line generated in this study, BT-474-Luc2 targeting by HER2 CAR-T cells could also be tested using the impedance assay. In this assay, as effector cells kill adherent cells and they detach from the plate, electrical resistance in the co-culture decreases (Lisby et al, 2021). BT-474-Luc2 cells alone or co-cultured with mock CAR-T cells at a 10:1 CAR-T to target cell ratio did not cause a change in cell impedance over time. However, when BT-474-Luc2 cells were co-cultured with HER2 CAR-T cells, impedance dramatically decreased, indicating detachment of BT-474-Luc2 cells from the plate (Fig 4B). Taken together, these results show that the BT-474-Luc2 line can be used to monitor HER2 CAR-T targeting efficacy either by luminescence or in real time by impedance.

### Using luminescence as a readout for ex vivo NK direct killing and ADCC

Another important area of cancer therapy research in addition to CAR-T therapy is the use of monoclonal antibodies (Lo Nigro et al., 2019). We next evaluated the ability of two of the luciferase lines, WIL2-S-Luc2 and BT-474-Luc2, to serve as reporters of antibody-dependent cellular cytotoxicity. We first conducted a direct killing assay using primary NK cells at varying NK to target cell ratios. After 16 hours of co-culture, luminescence dramatically decreased with increasing ratios of NK to target cells, indicating dose-dependent reporter cell killing by NK cells (Figs 4C and 4D). Using a NK to target cell ratio of 2:1, we next performed the luciferase assay with anti-CD20 and anti-HER2 antibodies for NK cell-mediated ADCC (antibody-dependent cell-mediated cytotoxicity) against Wil2-S-Luc2 and BT-474-Luc2 cells, respectively. Rituximab (anti-CD20) or Trastuzumab (anti-HER2) monoclonal antibodies were administrated at various concentrations using IgG1 as an isotype control. We observed a dose-dependent decrease in luciferase signal relative to IgG1-treated controls in WIL2-S-Luc2 and BT-474-Luc2 cells, indicating antibody-dependent cell killing (Figs 4E and 4F). This result demonstrates that these reporter lines can also be targeted by monoclonal antibodies and thus can be used to measure antibody-mediated cellular cytotoxicity.

### Optimizing ex vivo CD20 CAR-T cell-mediated cytotoxicity assay using luminescence and Cytotox Red dye based live cell imaging

Finally, we investigated whether the engineered Farage-Luc2 and Daudi-Luc2 lines could serve as reporters of CD20 CAR-T targeting. CD20-positive Farage-Luc2 cells or Daudi-Luc2 cells were co-cultured with either CD20 CAR-T or mock CAR-T cells at various ratios of CAR-T cells to target cells (1:1, 2:1, 5:1, and 10:1). After 24 hours, luminescence significantly decreased following co-culture with CD20 CAR-T cells relative to mock CAR-T controls as seen for the other luciferase lines (Figs 5A and 6A). We then used a tractable live imaging approach to assess cell death of the reporter lines after co-culture with CD20 CAR-T cells. Cytotox red, which only stains dead cells (Schwietzer et al., 2023), was added to co-cultures of each reporter line with either CD20 CAR-T cells or mock CAR-T cells at various ratios of CAR-T to target cells. Red fluorescence was then quantified after 24 hours of co-culture. While there was a slight increase in red fluorescence after co-culture with mock CAR-T cells, a dramatic and dose-dependent increase in red fluorescence intensity was observed after co-culture with CD20 CAR-T cells for both Farage-Luc2 and Daudi-Luc2 (Figs 5B and 6B). Live cell imaging of the co-cultures revealed a significant increase in red fluorescence over time in Farage-Luc2 or Daudi-Luc2 cells co-cultured with CD20 CAR-T cells compared to reporter cells co-cultured with mock CAR-T cells (Fig 5C and D; Fig 6C and D; Videos 3-6). These results indicate elevated levels of targeted cell killing of both reporter lines specifically by CD20 CAR-T cells and demonstrate another live imaging approach that can be used in parallel with luminescence.

**Figure 5:**
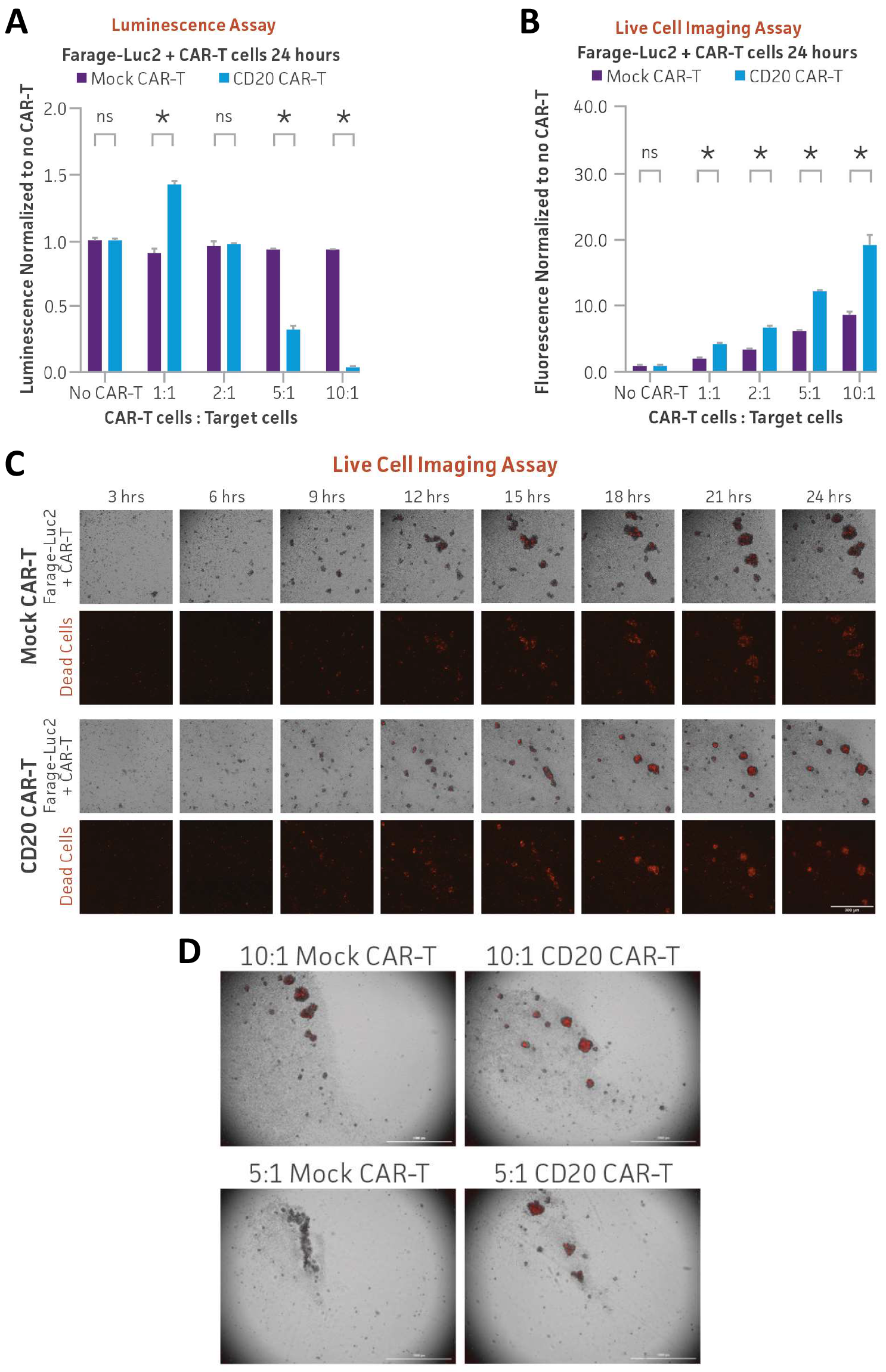
CD20 CAR-T in vitro killing assay of Farage-Luc2 measured using luminescence and live cell imaging. (A) CD20-positive Farage-Luc2 cells (5 x 10^3^) were seeded into a 96-well plate and CD20 CAR-T or mock CAR-T cells were added at various ratios of CAR-T to target cells (1:1, 2:1, 5:1, and 10:1). B) Farage-Luc2 cells (5 x 10^3^) were co-cultured with CD20 CAR-T cells or mock CAR-T cells in the presence of Incucyte Cytotox red dye. Fluorescent images were captured every hour for 24 hours and red fluorescence was quantified and plotted. * = significant difference and ns = not significant using an unpaired t test, with a single pooled variance. N=3 in all experiments. *P < 0.05. (C) Series of images captured during Farage-Luc2 cells co-culture with CD20 or mock CAR-T cells in the presence of Cytotox Red. Scale bar, 300 μm. (D) The larger fields of view of Farage-Luc2 cells co-culture with CD20 or mock CAR-T cells in the presence of Cytotox Red at 24 hours. Scale bar, 1000 μm.

**Figure 6:**
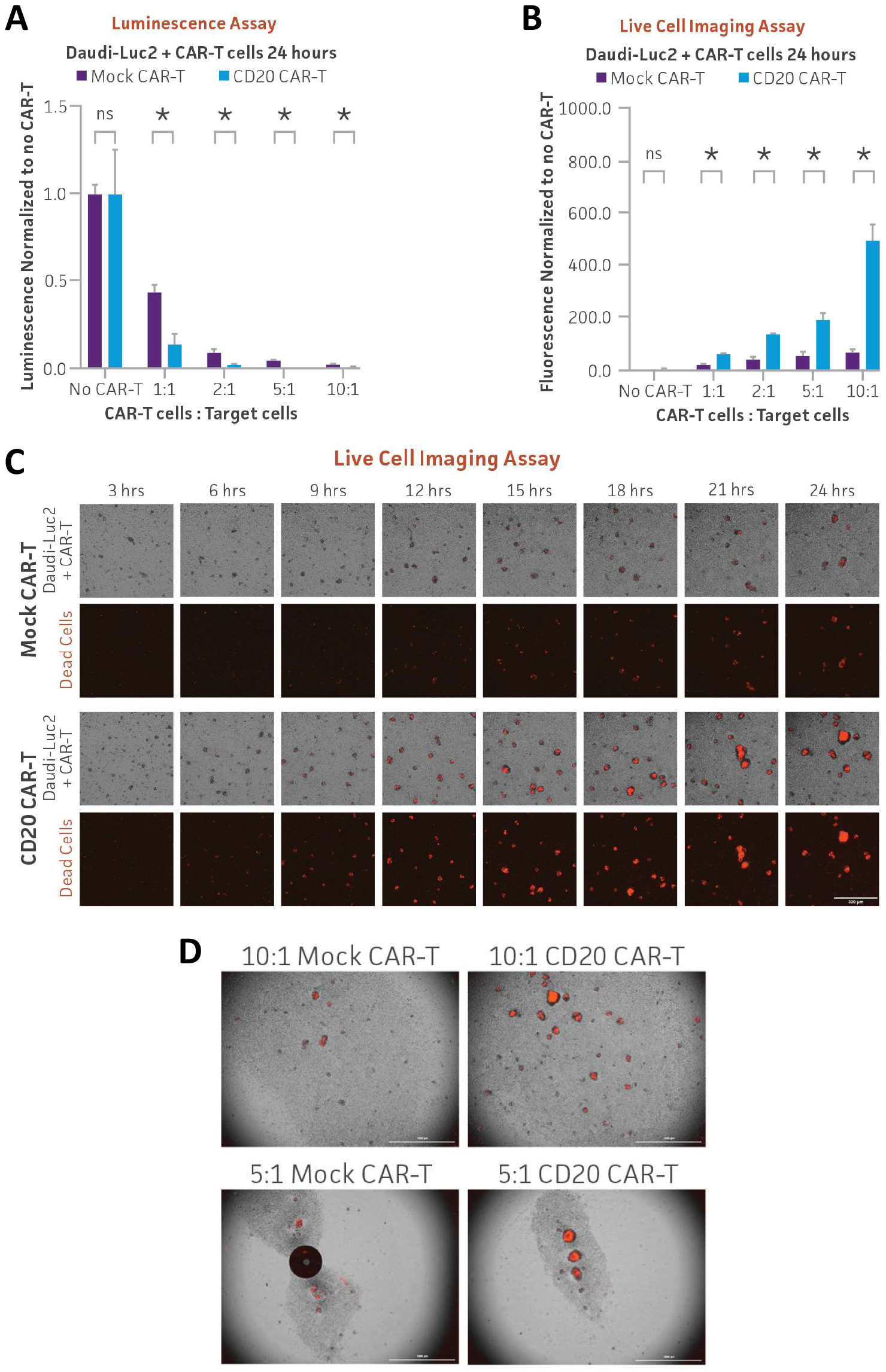
CD20 CAR-T in vitro killing assay of Daudi-Luc2 measured using luminescence and live cell imaging. (A) CD20-positive Daudi-Luc2 cells (5 x 10^3^) were seeded into a 96-well plate and CD20 CAR-T or mock CAR-T cells were added at various ratios of CAR-T cells to target cells (1:1, 2:1, 5:1, and 10:1). After 24 hours of co-culture, Bright-Glo was added to the wells and luminescence was measured. B) Daudi-Luc2 cells (5 x 10^3^) were co-cultured with CD20 CAR-T cells or mock CAR-T cells in the presence of Incucyte Cytotox red dye. Fluorescent images were captured every hour for 24 hours, and red fluorescence was quantified and plotted. * = significant difference and ns = not significant using an unpaired t test, with a single pooled variance. N=3 in all experiments. *P < 0.05. (C) Series of images captured during Daudi-Luc2 cells co-culture with CD20 or mock CAR-T cells in the presence of Cytotox Red. Scale bar, 300 μm. (D) The larger fields of view of Daudi-Luc2 cells co-culture with CD20 or mock CAR-T cells in the presence of Cytotox Red at 24 hours. Scale bar, 1000 μm.

## Discussion

In this study, we generated five luciferase reporter cancer cell lines and demonstrated their use in a variety of multimodality imaging approaches to measure CAR-T cell cytotoxicity. The CAR-T target luciferase reporter tumor cell lines that we engineered were derived from a variety of highly malignant liquid and solid cancer types, namely B cell lymphoma, Burkitt’s lymphoma, Non-Hodgkin’s B cell lymphoma and ductal breast carcinoma. These novel target cell lines were generated from parental tumor cells that have high endogenous expression of target antigens such as CD19, CD20, and HER2. Stable luciferase expressing clones were engineered to display high signal-to-noise ratios, aiding in interpretation of the data. Furthermore, authenticating these cell lines using tried and true methods such as short tandem repeat (STR) profiling, mycoplasma detection, cell growth rate and morphology assays can satisfy the requirements for authentication set by regulatory bodies and give researchers further confidence in their experimental results. The performance of the CAR-T target luciferase reporter cell lines was verified in T cell co-culture experiments. Commercially available CAR-T cells targeting CD19, CD20, and HER2 were employed in this study, with which non-targeting mock CAR-T cells from the same donor were paired as controls. The cytotoxicity of the CAR-T cells against target tumor cells was measured using a luciferase assay, a commercially available impedance assay, and two different fluorescent live imaging assays. Our results demonstrate that the luciferase reporter system is a simple, robust, and highly sensitive means to measure biological processes in cancer and T cell ex vivo co-cultures through a variety of multimodality imaging methods.

Assays currently in use have various drawbacks (Table 1). Unlike the luminescence and live imaging assays, the chromium release assay uses radioactive isotopes that can pose safety concerns and requires specialized handling and disposal procedures. The assay also just provides a single-time-point measurement and does not allow for real-time or kinetic assessment of CAR-T cell cytotoxicity over time (Lisby et al., 2022; Peper et al., 2014) like live imaging allows. Some background release of chromium can occur even in the absence of CAR-T cells, leading to false-positive results and potentially affecting the accuracy of the assay (Kiesgen et al., 2021; Peper et al., 2014; Riss et al., 2019). Further, the chromium release assay may not be sensitive enough to detect low levels of target cell killing, especially when CAR-T cells are present in limited numbers or exhibit lower cytotoxic activity (Kiesgen et al., 2021). As demonstrated previously, the luciferase assay is highly sensitive (Matta et al., 2018; Schäfer et al., 1997). We further show here that a reduction in luminescence can be detected even at low ratios of effector to target cells. The chromium release assay requires physical contact between effector CAR-T cells and target cells to measure cytotoxicity accurately. However, certain CAR-T cell constructs or target cells may have limited adhesion capabilities or poor compatibility, leading to reduced accuracy or underestimation of cytotoxicity. Moreover, the chromium release assay requires time for the released chromium to be measured, which introduces a delay between target cell lysis and data acquisition (Peper et al., 2014). This delay may not capture the full kinetics of CAR-T cell cytotoxicity and can impact the accuracy of time-sensitive measurements. The luciferase assay avoids this delay before lysis, as data is acquired within minutes.

**Table 1:** Comparison of cell-mediated cytotoxicity assays.

|  | Chromium release | Bioluminescence imaging | Impedance | Flow cytometry | Fluorescence Imaging - cell labeling | Fluorescence imaging - cytotox dye in media |
| --- | --- | --- | --- | --- | --- | --- |
| Principal measure of cytotoxicity | <sup>51</sup>Cr release | Luciferase activity | Cell detachment | Live/dead staining phenotype | Decrease in fluorescent signal | Dye infiltration into dead cell |
| Radioactive materials needed | Yes | No | No | No | No | No |
| Target cell labeling required | Yes | No | No | Yes | Yes | No |
| Genetic modification of target cells | No | Yes (reporter gene) | No | No | No | No |
| Endpoint/kinetic | Endpoint | Temporal & Spatial | Temporal | Endpoint | Temporal & Spatial | Temporal & Spatial |
| Real-time measurement | No | Yes | Yes | No | Yes | Yes |
| Maximum time point measured | 18 – 24 hours | Days | Days | Days | Days | Days |
| Ability to measure different cytotoxicity heterogenous targets | No | No | No | Yes | Yes | Yes |
| Throughput and automatability | Low | High | High | High | High | High |

Impedance-based assays typically measure overall changes in electrical impedance and cannot distinguish between CAR-T cell-mediated cytotoxicity and non-specific effects, such as changes in cell adherence or morphology, which live imaging can capture. Impedance assays may lack the sensitivity to detect low levels of target cell killing, especially when CAR-T cells are present in small numbers or have lower cytotoxic potential. As mentioned above, the luciferase assay can detect a decrease in signal even at low effector to target cell ratios. Impedance-based assays are often limited to measuring a single parameter at a time and may not easily allow for multiplexing or simultaneous evaluation of multiple aspects of CAR-T cell cytotoxicity. Furthermore, impedance-based assays measure changes in electrical impedance caused by cellular interactions but do not directly measure cytotoxicity (Witzel et al., 2015). Changes in impedance may be influenced by factors other than cytotoxicity, such as changes in cell adherence, morphology, or cell viability (Giaever and Keese, 1993). This lack of direct measurement of cytotoxicity can introduce some degree of uncertainty and affect the accuracy of the assay. While both the luciferase and live imaging assays require labeling, any changes in luminescence or fluorescence are a direct result of cell death. Further, impedance-based assays may have a limited ability to detect low levels of target cell killing or discriminate between different degrees of cytotoxicity. This limitation can lead to underestimation of CAR-T cell activity when cytotoxicity is low or subtle, which is more likely to be captured by the quantitative sensitivity of the luciferase assay and continual monitoring by live imaging.

Lastly, flow cytometry assays may have a limited dynamic range, making it challenging to accurately measure high levels of target cell killing or discriminate between different degrees of cytotoxicity (Langhans et al., 2005). Flow cytometry provides information on individual cells but does not provide spatial information about the interactions between CAR-T cells and target cells within the tumor microenvironment (Carannante et al., 2023). A major strength of the live imaging assay is the ability to capture these interactions. Non-specific binding of antibodies or detection reagents to cells can introduce background noise, potentially impacting the accuracy of the assay and leading to false-positive or false-negative results. Furthermore, flow cytometry assays rely on fluorophore-labeled antibodies for target cell detection. The choice of fluorophores can impact the accuracy and sensitivity of the assay. Some fluorophores may have limited brightness, photobleaching, or spectral overlap, leading to decreased detection sensitivity or potential interference with other fluorescence channels (Flores-Montero et al., 2019). Non-specific binding of antibodies or detection reagents to cells can generate false-positive results, particularly when dealing with low-level antigen expression on non-target cells. This non-specific binding can result in overestimation of CAR-T cell cytotoxicity or false identification of target cells. While the luciferase assay necessitates genetic modification and live imaging requires labeling via dyes, these labels are highly specific to the target cells, avoiding many of these drawbacks. Moreover, flow cytometry assays primarily focus on surface marker expression or cell death markers, but they may not capture other phenotypic changes or functional alterations in CAR-T cells that could affect their cytotoxic potential.

Overall, the luminescence assay’s quantitative power, coupled with kinetic insights from live imaging, renders these multimodality imaging assays highly efficient for real-time monitoring of ex vivo CAR-T cell-mediated cytotoxicity. While each CAR-T cytotoxicity assay has inherent limitations, utilizing luciferase reporter CAR-T target tumor cell lines in combined multimodality imaging assays creates an optimized platform to facilitate the collection of more comprehensive cell information during CAR-T-mediated tumor cell elimination. The generated lines, along with other ATCC luciferase tumor lines expressing key CAR-T targets, serve as valuable and reliable tools for evaluating CAR-T targeting efficacy in the advancement of enhanced CAR-T therapies.

## Materials and Methods

### Cell culture, lentiviral transfection, and single cell cloning

The following cell lines were obtained from ATCC based upon their high expression levels of common CAR-T target antigens and grown according to the ATCC product sheet recommendations: WIL2-S, Raji, Daudi, Farage, and BT-474. 500,000 cells per line were transduced with a lentiviral plasmid expressing luciferase under the control of an EF1A promoter using 50 μg/mL of protamine sulfate. After viral transduction, Luc2 cell lines were selected and maintained in the same culture media as the parental cell lines but with 4.0 – 8.0 μg/mL blasticidin (Gibco, Waltham, MA). After selection, luciferase pools were screened for luciferase activity. Single cells were sorted into 96-well plates using a Sony SH800 automated cell sorter (Sony Biotechnology, San Jose, CA) and expanded for approximately 10-14 days until confluency reached 70%. The single clones were then subcultured and evaluated for luciferase activity again. The single clones that yielded the highest luciferase activity were chosen for further evaluation to verify that the introduction of luciferase into the cell line did not cause changes in growth rate, morphology, and expression of target antigens.

### Next generation sequencing and expression heat map generation

RNA sequencing was performed in duplicate for Raji and WIL2-S cell lines and their paired Luc2 daughter lines. Library preparation using the TruSeq Stranded mRNA library prep kit (Illumina, San Diego, CA) and 150 bp paired-end sequencing on a NovaSeq6000 S4 were performed at Psomagen (Rockville, MD). Then reads were aligned to hg38 using HISAT2 (v2.1.0) and transcripts were assembled using StringTie (v2.1.3b) at Psomagen. Expression heat maps were generated using Microsoft Excel for 12 hematologic tumor CAR-T targets and 26 solid tumor CAR-T targets either using the Cancer Cell Line Encyclopedia (CCLE) database (Broad Institute, Cambridge, MA) or the in-house bulk RNA-seq results. The CCLE values are log2(TPM+1) according to the CCLE public data guidance. The in-house RNA-seq values are log2(FPKM+1).

### Co-culture setup for CAR-T cytotoxicity assays

CAR-T cytotoxicity was studied using CD19 CAR-T, CD20 CAR-T, or HER2 CAR-T cells (ProMab, Richmond, CA) which were each paired with mock CAR-T cells generated from the same donor as a control. CAR-T target luciferase reporter cell lines were seeded at 5 x 10^3^ cells per well into 96-well plates (Corning, Corning, NY) in triplicate in 50 μl of complete media (without blasticidin). 50 μl of CAR-T cells were added to each well to correspond to an effector to target ratio (CAR-T cells: Target cells) of 1:1, 2:1, 5:1 or 10:1. The CAR-T target luciferase reporter cells were either pre-stained with Vybrant™ DiO (Thermo Fisher, Waltham, MA), co-cultured in the presence of Incucyte® Cytotox red dye (Sartorius, Göttingen, Germany), or left unstained for the luciferase assay.

### Luminescence assay

CAR-T cytotoxicity co-culture was performed as previously described. After 24 hours of co-culturing, an equal volume of Bright-Glo™ Reagent (Promega, Madison, WI) was added to the cells. After 8 minutes of incubation at room temperature on a plate shaker, reactions were transferred to a white opaque plate and luminescence was measured on a SpectraMax i3x plate reader (Molecular Devices, San Jose, CA). Luminescence was normalized to wells with no CAR-T cells and was plotted as fold change in Prism 9 v9.0.0 (121) (Graphpad, San Diego, CA).

### Real-time xCELLigence live cell analysis

BT-474-Luc2 cells (2 x 10^4^ cells/well) were seeded in 96-well RTCA E-Plate (Agilent, Santa Clara, CA) and left to adhere for 24 hours. On the day of the co-culture cytotoxicity experiments, HER2 CAR-T cells or mock CAR-T cells from the same donor were added at an effector:target cell ratio of 10:1 and measured in a real-time cell analysis assay using the xCELLigence SP system (Agilent, Santa Clara, CA). An electrical current was applied to each co-culture. E-plates were read every 15 minutes for 24 hours and data were normalized to starting resistance. The normalized cell index was plotted as fold change against time in Prism 9 v9.0.0 (121) (Graphpad, San Diego, CA).

### DiO staining for live cell imaging

5 μl of Vybrant DiO cell labeling solution (Molecular Probes, Eugene, OR) was added to 1 mL of serum-free RPMI-1640 media (ATCC, Manassas, VA), premixed by vortexing, and used to resuspend Raji-Luc2 cells at 1 x 10^6^ cells/ml. Cells were transferred to a 12-well plate and incubated at 37 °C for 45 minutes with agitation every 5-10 minutes. Cells were pelleted by centrifugation and washed 3x with fresh culture media. Cells were plated and allowed to recover overnight prior to seeding into a 96-well plate for co-culture experiments. After the co-culture assay was set up, green fluorescent and brightfield images were captured with the Cytation1 plate reader (Agilent, Santa Clara, CA) using a 4x objective to capture 2 x 2 image montages in 30-min intervals for 24 hours. Images were stitched using Gen 5 v3.11 software (Agilent, Santa Clara, CA).

### Cytotox Red dye staining for live cell imaging

CAR-T cytotoxicity co-culture was set up as previously mentioned into 96 well plates. Cytotox Red (Sartorius, Göttingen, Germany) was diluted to a stock of 100 uM using DPBS and then was used at a final concentration of 250 nM in the well. Red fluorescent and brightfield images were captured with the Cytation1 plate reader (Agilent, Santa Clara, CA) using a 4x objective to capture 2 x 2 image montages in 1-hour intervals for 24 hours. Images were stitched using Gen 5 v3.11 software (Agilent, Santa Clara, CA). Red fluorescent intensity was either measured using Gen5 software or ImageJ v1.53t (NIH, Bethesda, MD).

### NK direct killing and ADCC assay

2 x 10^6^ naive NK cells (ATCC, Manassas, VA) were stimulated with MACS beads (Miltenyi Biotech, Bergisch Gladbach, Germany) and cultured for 7 days for expansion. Luciferase-expressing cells were seeded at 5.0 x 10^3^ cells/well in triplicate into 96-well plates using RPMI+10% low IgG FBS medium. Various ratios of target cells to effector cells ranging from 1:4 to 16:1 was tested for direct cell killing via luminescence. To perform the ADCC assay, a 96-well plate was set up with Rituximab (anti-CD20), Trastuzumab (anti-HER2), or isotype control mAb antibody (Invivogen, San Diego, CA) at concentrations from 10 μg/mL diluting down to 0.0001 μg/mL. 5 x 10^3^ target cells and 2.5 x 10^3^ NK cells were added for a 2:1 target to effector cell ratio. Co-cultures were incubated for 4 hours. Then 100 μl of the co-culture was transferred to a white plate with an equal volume of Bright-Glo reagent (Promega, Madison, WI) and was incubated for 5 minutes before luminescence was measured.

### Flow cytometry

For the flow cytometry analyses, cells were stained with the following fluorescently labeled antibodies specific for CAR-T target antigens on the cell surface: APC anti-CD19, FITC anti-CD20, and PE anti-HER2 (Miltenyi Biotec, Bergisch Gladbach, Germany). PE, APC, or FITC-conjugated human IgG1 (Miltenyi Biotec) was used as an isotype control. Cell staining was performed in Fetal Bovine Serum Stain Buffer (BD Biosciences™) with the antibodies diluted to the manufacturers’ recommended concentrations for 30 minutes on ice in the dark. Cells were then fixed by the addition of an equal volume of Fixation Buffer (BD Biosciences™). For each sample, 10,000 stained cells were captured by the Accuri C6 Plus flow cytometer (BD) and the results were analyzed by the FlowJo® software version v10.9.0.

## Data availability

The data presented in this study are available on request from the corresponding author.

## Supplementary Information

Supplementary information is attached in a separate pdf for submission.

## Acknowledgements

This research received no external funding. We thank Brian Shapiro and Cara Wilder for helping edit the manuscript. We thank Andrew Burke for optimizing the figures and graphs.

## Author Contributions

Conceptualized the project and designed the methodology, J.G.F, F.T., and Z.G.; Performed experiments, analyzed data, and contributed to writing the manuscript, J.G.F, L.C., H.C. and C.E.M.; Supervised all aspects of the project and wrote the manuscript, F.T. and Z.G.; All authors have read and agreed to the published version of the manuscript.

