## Supplementary Figures for "Optimizing ex vivo CAR-T cell-mediated cytotoxicity assay through multimodality imaging"

Methods

#### Supplementary Figure and Video Legends

**Figure S1: Characterization of CAR-T target cell lines: Cell Morphology.** Cell morphology of the luciferase-expressing cell lines compared to their parental lines. Scale bar, 100 µm.

**Figure S2: Characterization of CAR-T target cell lines: Growth Curve.** 2 x $10^5$ cells/mL were seeded into T25 flasks and cell counts were performed every day for 5, 7 or 9 days. The parental cell lines are shown in black, and the CAR-T target luciferase reporter cell lines are shown in green.

**Figure S3 Characterization of CAR-T target cell lines: Antigen FACS analysis.** Flow cytometry was performed to assess the CAR-T target antigen expression levels of CD19, CD20, and HER2 (pink) on the tumor cell lines compared to isotype controls (blue).

**Figure S4: Characterization of CAR-T target cell lines: Luciferase assay.** Luciferase expression was measured using Bright-Glo™ reagent and a luminescence plate reader for increasing numbers of cells. The results show a linear correlation between bioluminescence intensity and cell number.

**Figure S5: Characterization of CAR-T target cell lines: Luciferase activity stability.** To verify the stability of luciferase expression, the cells were maintained in culture for 30 population doublings. Luciferase expression was monitored every week.

**Videos 1 and 2: CD19 CAR-T in vitro killing assay of Raji-Luc2 measured using live cell imaging.** Raji-Luc2 cells were stained with Vybrant DiO dye and real-time fluorescent images were captured every 30 minutes for 24 hours during co-culture with either CD19 or mock CAR-T cells.

**Videos 3 and 4: CD20 CAR-T in vitro killing assay of Farage-Luc2 measured using live cell imaging.** Farage-Luc2 cells co-culture with CD20 or mock CAR-T cells in the presence of Incucyte Cytotox red dye. Real-time fluorescent images were captured every 1 hour for 24 hours during the co-culture. Scale bar, 1000 µm.

**Videos 5 and 6: CD20 CAR-T in vitro killing assay of Daudi-Luc2 measured using live cell imaging.** Daudi-Luc2 cells co-culture with CD20 or mock CAR-T cells in the presence of Incucyte Cytotox red dye. Real-time fluorescent images were captured every 1 hour for 24 hours during the co-culture. Scale bar, 1000 µm.

### Supplemental Fig. S1

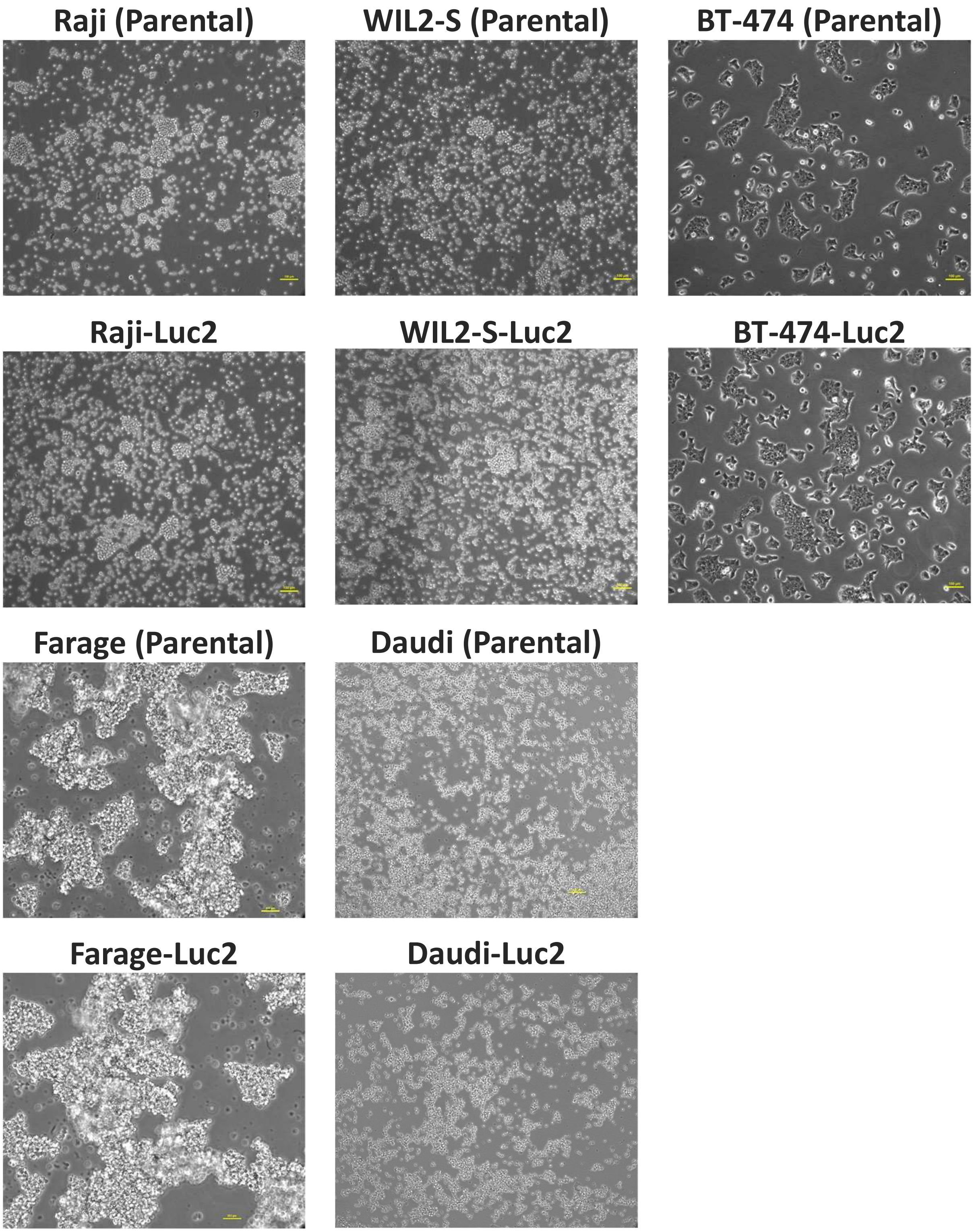


**Figure S1: Characterization of CAR-T target cell lines: Cell Morphology.** Cell morphology of the luciferase-expressing cell lines compared to their parental lines. Scale bar, 100 µm.

### Supplemental Fig. S2

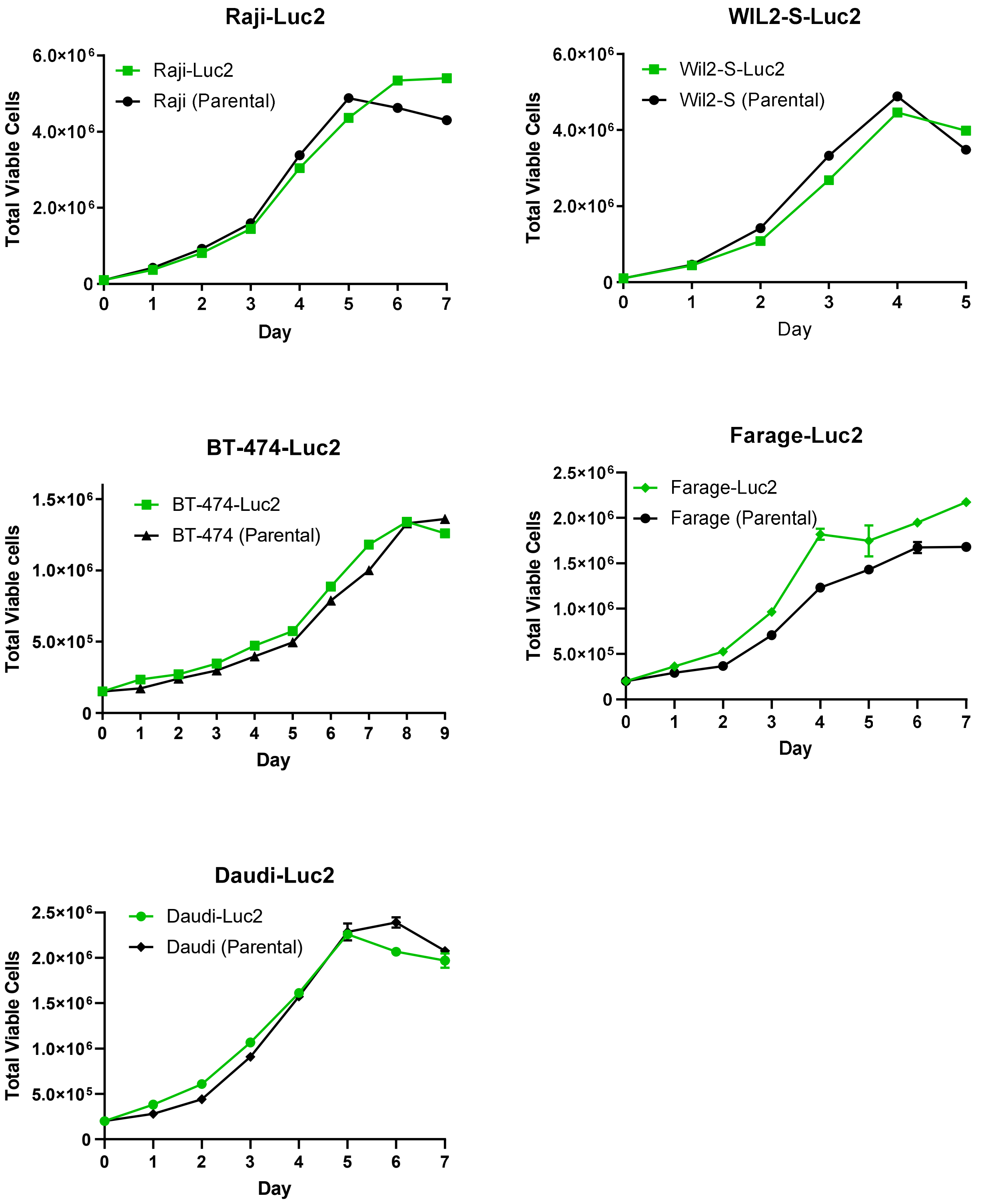


**Figure S2: Characterization of CAR-T target cell lines: Growth Curve.** 2 x $10^5$ cells/mL were seeded into T25 flasks and cell counts were performed every day for 5, 7 or 9 days. The parental cell lines are shown in black, and the CAR-T target luciferase reporter cell lines are shown in green.

#### Supplemental Fig. S3

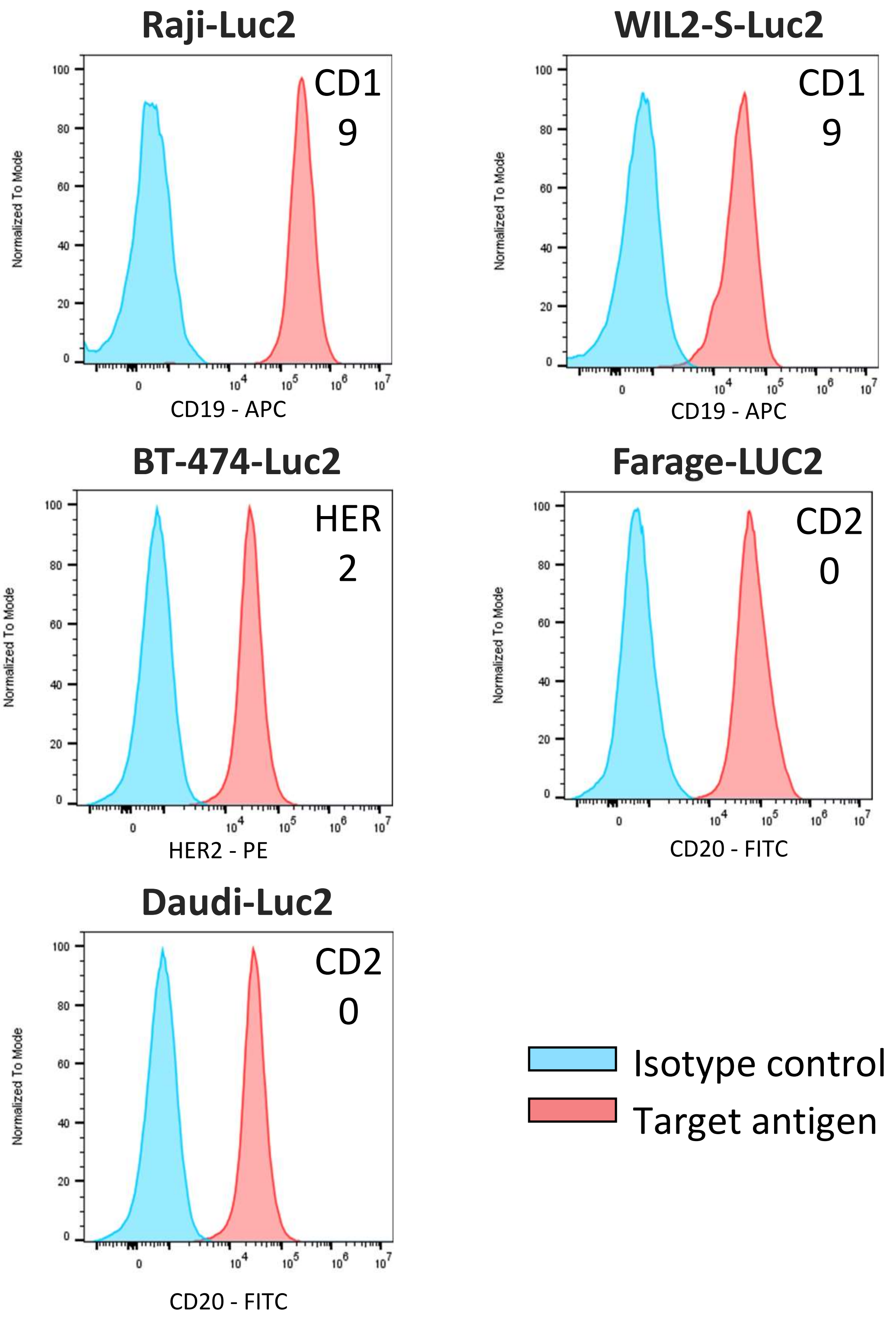


**Figure S3 Characterization of CAR-T target cell lines: Antigen FACS analysis.** Flow cytometry was performed to assess the CAR-T target antigen expression levels of CD19, CD20, and HER2 (pink) on the tumor cell lines compared to isotype controls (blue).

### Supplemental Fig. S4

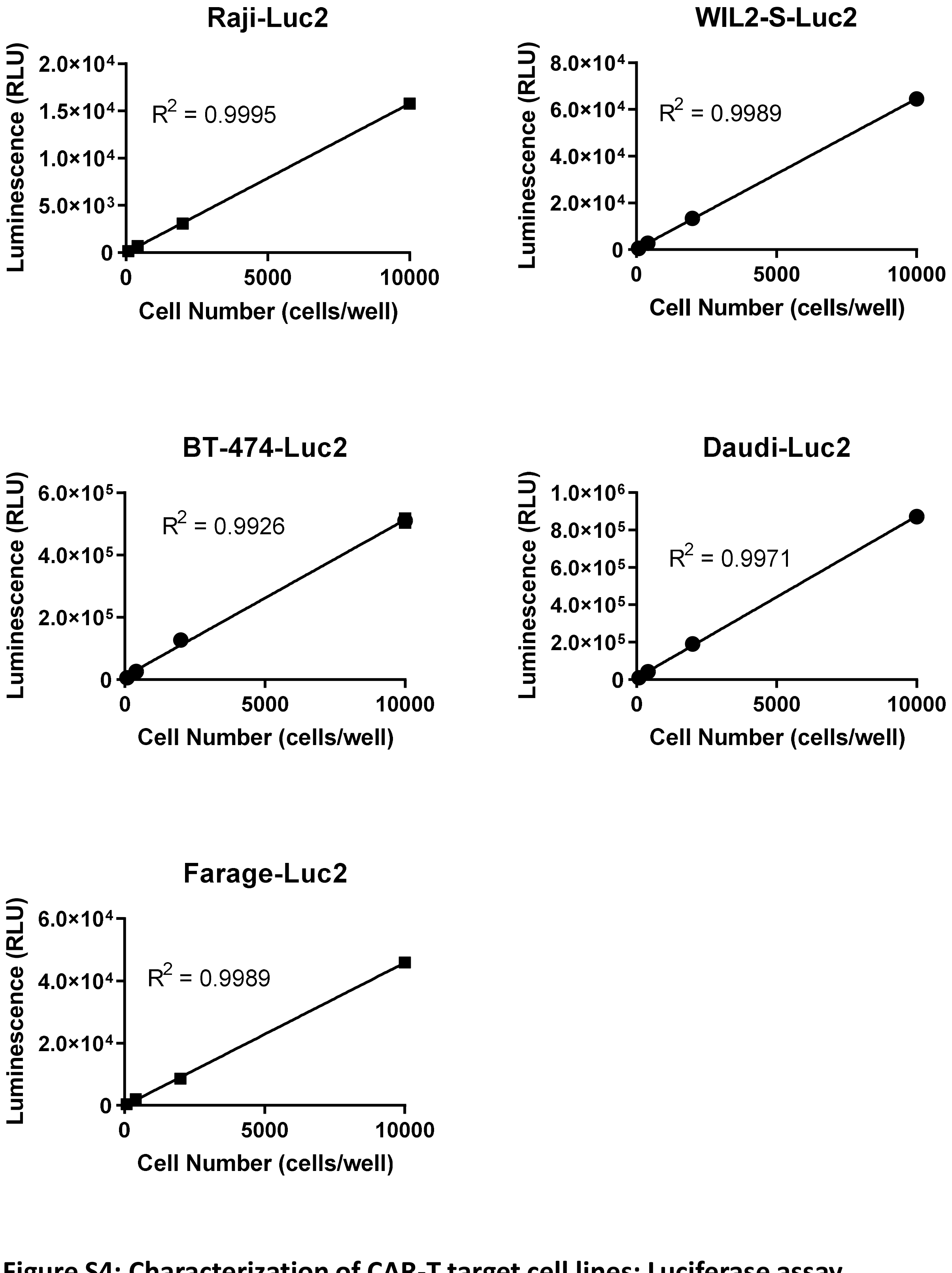


**Figure S4: Characterization of CAR-T target cell lines: Luciferase assay.** Luciferase expression was measured using Bright-Glo™ reagent and a luminescence plate reader for increasing numbers of cells. The results show a linear correlation between bioluminescence intensity and cell number.

### Supplemental Fig. S5

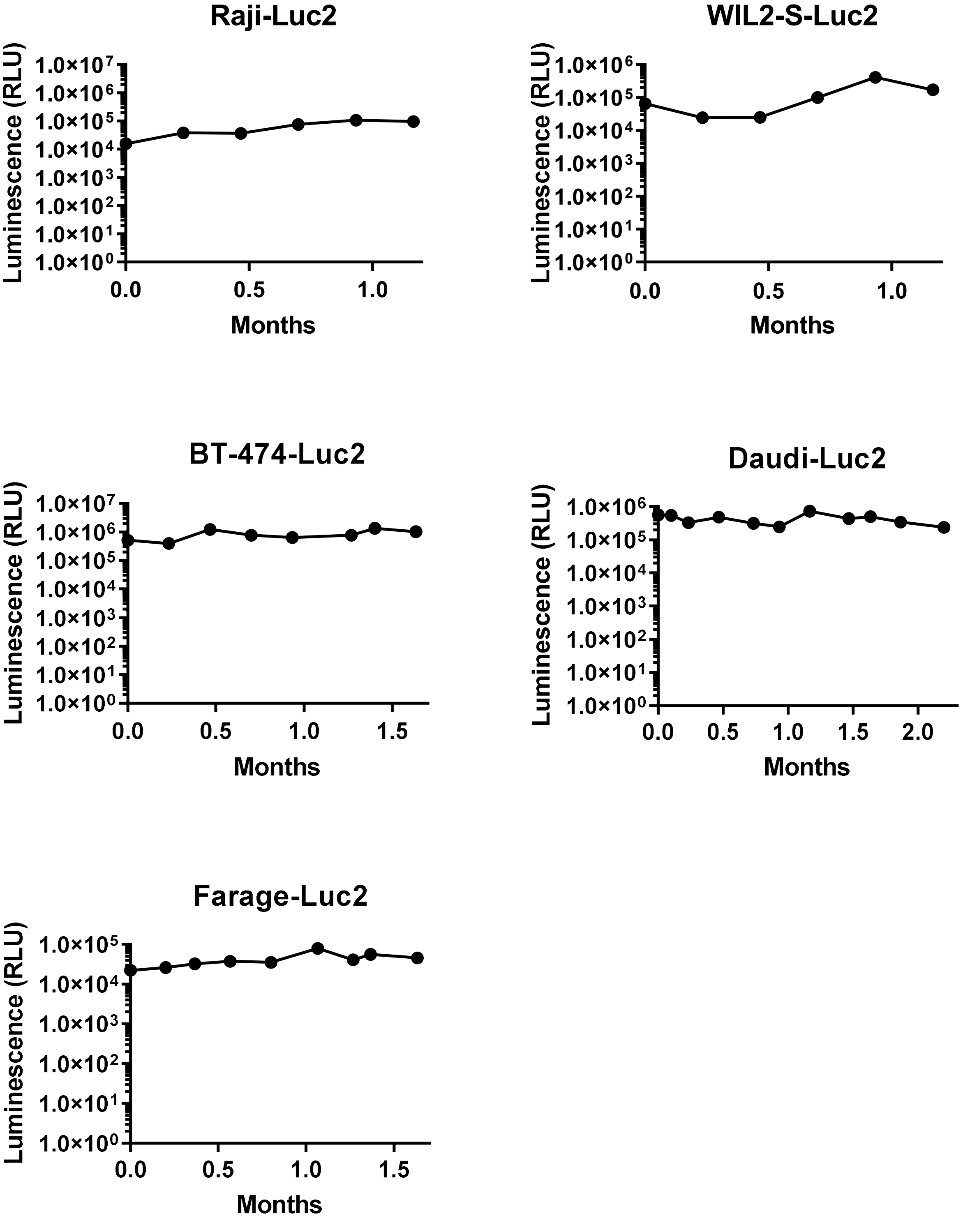


**Figure S5: Characterization of CAR-T target cell lines: Luciferase activity stability.** To verify the stability of luciferase expression, the cells were maintained in culture for 30 population doublings. Luciferase expression was monitored every week.
